# The Human Fragile X Mental Retardation Protein Inhibits the Elongation Step of Translation through its RGG and C-terminal domains

**DOI:** 10.1101/2020.06.25.171967

**Authors:** Youssi M. Athar, Simpson Joseph

## Abstract

Fragile X mental retardation protein (FMRP) is an RNA-binding protein that regulates the translation of numerous mRNAs in neurons. The precise mechanism of translational regulation by FMRP is unknown. Some studies have indicated that FMRP inhibits the initiation step of translation, whereas other studies have indicated that the elongation step of translation is inhibited by FMRP. To determine whether FMRP inhibits the initiation or the elongation step of protein synthesis, we investigated m^7^G-cap-dependent and IRES-driven, cap-independent translation of several reporter mRNAs in vitro. Our results show that FMRP inhibits both m^7^G-cap-dependent and cap-independent translation to similar degrees, indicating that the elongation step of translation is inhibited by FMRP. Additionally, we dissected the RNA-binding domains of hFMRP to determine the essential domains for inhibiting translation. We show that the RGG domain, together with the C-terminal domain (CTD), is sufficient to inhibit translation while the KH domains do not inhibit mRNA translation. However, the region between the RGG domain and the KH2 domain may contribute as NT-hFMRP shows more potent inhibition than the RGG-CTD tail alone. Interestingly, we see a correlation between ribosome binding and translation inhibition, suggesting the RGG-CTD tail of hFMRP may anchor FMRP to the ribosome during translation inhibition.

## Introduction

Fragile X syndrome (FXS) is the most common form of inherited intellectual disability and is caused by the reduced expression of the fragile X mental retardation 1 (*Fmr1*) gene in neurons [1]. The expression of the *Fmr1* gene is reduced because of the expansion of the CGG trinucleotide repeats in the 5’-untranslated region of the gene, which results in abnormal methylation of the gene and the repression of transcription [2–5]. *Fmr1* encodes an RNA binding protein, fragile X mental retardation protein (FMRP), that is highly expressed in the brain [6–10]. FMRP has four RNA-binding domains: one RGG domain that is rich in arginines and glycines and three hnRNP K-homology domains (KH0, KH1, and KH2) **(Figure 1)** [6,7,11]. FMRP is thought to regulate the translation of about 4% of the human fetal brain mRNAs by directly binding to specific sequences or structures in the target mRNAs [12]. Additionally, the majority of FMRP in the cell is associated with polyribosomes, consistent with its proposed role in regulating protein synthesis [13]. Interestingly, missense mutations in the KH1 and KH2 domains (Gly266Glu and Ile304Asn, respectively, of human FMRP) abolishes the binding of FMRP to polyribosomes and cause FXS in humans [14,15]. These results suggest that the RNA-binding activity of FMRP is critical for the healthy development and function of the brain.

**Figure 1.**
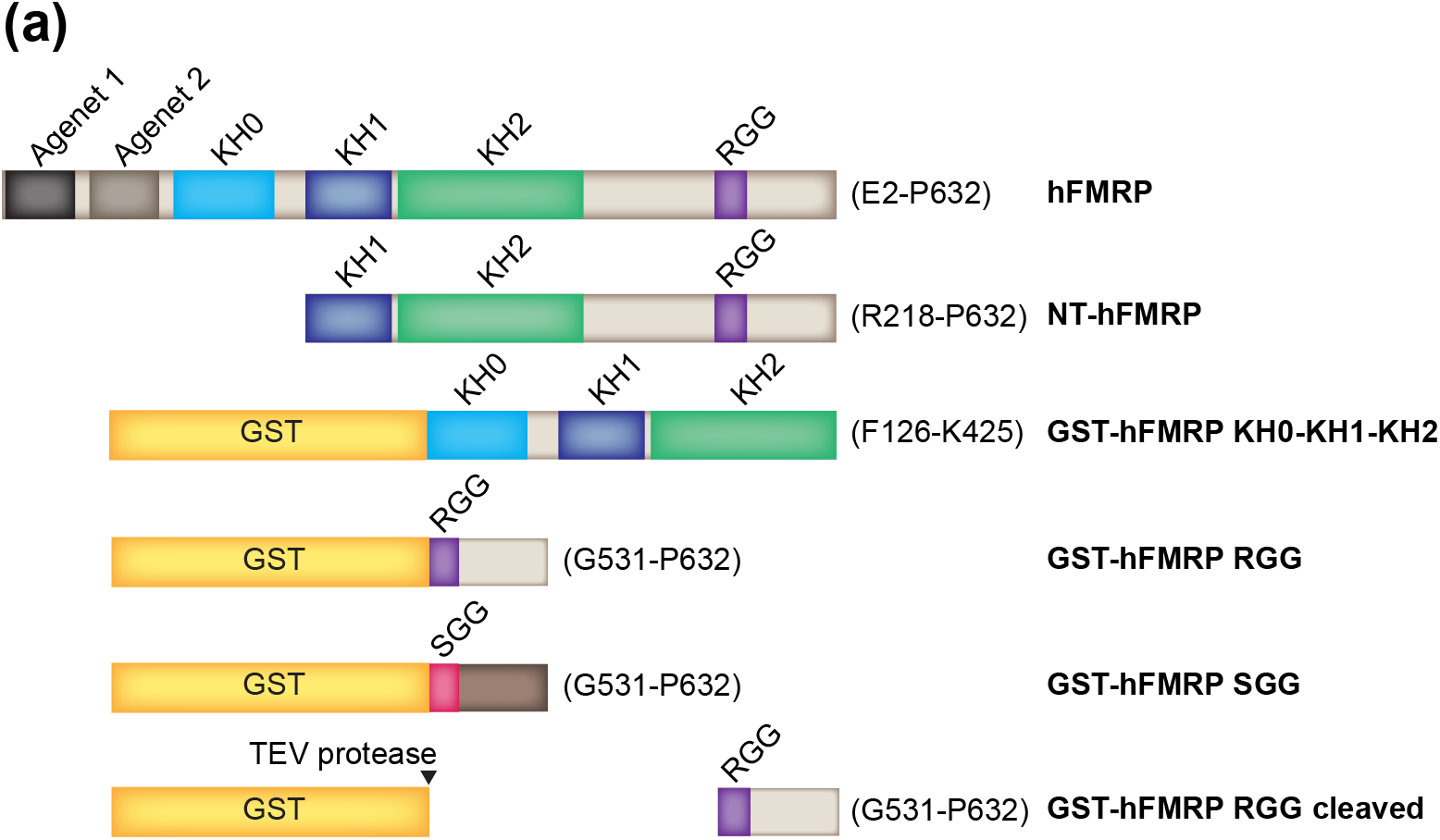
Human FMRP constructs. hFMRP is the full-length human FMRP isoform 1, spanning E2-P632, alongside the truncation constructs that were used. GST-hFMRP SGG is a fusion between the glutathione S-transferase and the RGG motif-containing sequence of hFMRP spanning G531-P632 where the 16 arginines spanning the RGG motif to the C-terminus were mutated to serines, illustrated using a different color for the C-terminus.

Several studies have focused their efforts on trying to identify the mRNA targets of FMRP. Although thousands of mRNA targets have been identified over the years, there is minimal overlap among the mRNA targets from individual studies [12,13,16–19]. These results suggest that it is difficult to discern the authentic mRNA targets of FMRP. Interestingly, in vitro selection experiments identified a G-quadruplex (GQ) structure and a pseudoknot structure as the potential RNA ligands for the RGG and KH2 domains, respectively [12,20–22]. Based on these results, it was proposed that FMRP may bind to mRNAs that possess GQ or pseudoknot forming sequences to repress their translation [12,20,22,23]. We previously showed that the KH domains of FMRP do not bind to simple four or five nucleotide RNA sequence motifs compared to the canonical KH domains found in other RNA-binding proteins [24–29]. It is possible that the KH domains of FMRP bind to pseudoknot structures in the mRNAs; however, such structures are not common in mRNAs and do not agree with the fact that FMRP appears to interact with thousands of mRNAs. In contrast, the RGG domain of FMRP binds to GQ structures with high affinity, and potential GQ structures are prevalent in mRNAs [24,30–36]. Interestingly, GQ structures are even present in the 28S and 18S rRNA expansion segments [37,38]. Therefore, possibly the RGG domain of FMRP interacts with GQ structures in the mRNA target or in the rRNAs to regulate translation.

The mechanism(s) by which FMRP regulates translation, however, is still unknown. One theory is that FMRP hinders translation initiation by binding to the target mRNA and recruiting the cytoplasmic FMRP-interacting protein (CYFIP1), which then binds the eukaryotic initiation factor 4E (eIF4E) at the mRNA 5’ 7-methylguanosine (m^7^G) cap. Binding of CYFIP1 to eIF4E, in turn, prevents binding of eukaryotic initiation factor 4G (eIF4G) to eIF4E, and the eukaryotic initiation factor 4F (eIF4F) complex is prevented from assembling at the m^7^G cap [39]. However, most of the cytoplasmic FMRP has been found associated with stalled polyribosomes [13,40]. Additionally, FMRP has been shown to interact with the ribosome even after translation initiation is blocked [41]. These results suggest that FMRP inhibits translation by binding to the target mRNA and stalling the ribosome during the translational elongation step.

Consistent with the idea that FMRP inhibits the elongation step, a cryo-electron microscopy (cryo-EM) structure of the *Drosophila* FMRP (dFMRP) bound to the *Drosophila* 80S ribosome showed that FMRP binds to a site within the ribosome that would prevent binding of tRNA and translation factors [42]. Additionally, the FMRP-ribosome structure revealed that the KH1 and KH2 domains bind near the ribosome’s peptidyl site, and the RGG domain of FMRP residing near the aminoacyl site, presumably available for binding target mRNA motifs. Thus, FMRP may tether itself to the ribosome and the mRNA via its KH and RGG domains, respectively, to stall the ribosome during the elongation step of translation [12,13,20,21,42,43].

FMRP is also suggested to repress translation via RNA interference (RNAi). FMRP has been shown to associate with the RNA-induced silencing complex (RISC) proteins Dicer and Argonaute 2 (Ago2) as well as specific microRNAs (miRNAs) [44–46]. In fact, FMRP has been reported to assemble the Ago2 and miRNA-125a inhibitory complex on the postsynaptic density protein 95 (PSD-95) mRNA through phosphorylation of FMRP [47,48]. The PSD-95 RNAi case may be just one of many RNAi translation repression pathways FMRP employs. It is possible that FMRP can recruit from a catalog of miRNA complexes to inhibit a variety of mRNAs via RNAi.

FMRP may use multiple mechanisms to specifically repress an extensive catalog of target mRNAs under different conditions within the cell. To study the mechanism of translational inhibition by FMRP, we analyzed the translation of 5’-m^7^G-capped mRNAs and uncapped mRNAs initiated using different IRES structures. Our studies show that FMRP reduces both m^7^G-cap-dependent and the cricket paralysis virus (CrPV) IRES-driven translation to a similar extent demonstrating that FMRP inhibits the elongation step of translation. Furthermore, we show that the RGG domain, together with the C-terminal domain (CTD), is sufficient to inhibit translation, whereas the KH domains do not inhibit mRNA translation. Interestingly, the RGG-CTD tail of FMRP binds to the ribosome suggesting that translation inhibition may depend on the interaction of FMRP with potential GQ structures in the rRNAs.

## Results

### FMRP inhibits the translation of various mRNAs

We sought to identify how the RNA-binding domains of human FMRP (hFMRP) contributed to the regulation of mRNA translation. We dissected hFMRP into two halves, the KH0-KH2 domains and the RGG domain with the C-terminal domain (CTD) **(Figure 1)**. We used a nuclease-treated rabbit reticulocyte lysate (RRL) in vitro translation (IVT) system to test FMRP effects on the *Renilla* luciferase (RLuc) reporter mRNA translation. First, we measured the importance of two canonical post-transcriptional modifications, the 5’-m^7^G cap and 3’ polyA tail, in increasing translation efficiency in the RRL system **(Figure 2)**. We observed that the individual modifications are not critical for efficient translation, but removing the 5’-m^7^G cap and 3’ polyA tail, significantly decreases translation efficiency in RRL **(Figure 2b)**. FMRP has been proposed to regulate the translation of some mRNAs during the initiation step through interactions involving the 5’ cap [39]. Therefore, we tested how NT-hFMRP regulates the translation of a polyadenylated RLuc with and without the 5’ cap. We observed FMRP inhibits both mRNAs similarly, suggesting that the inhibition is independent of the 5’ cap machinery.

**Figure 2.**
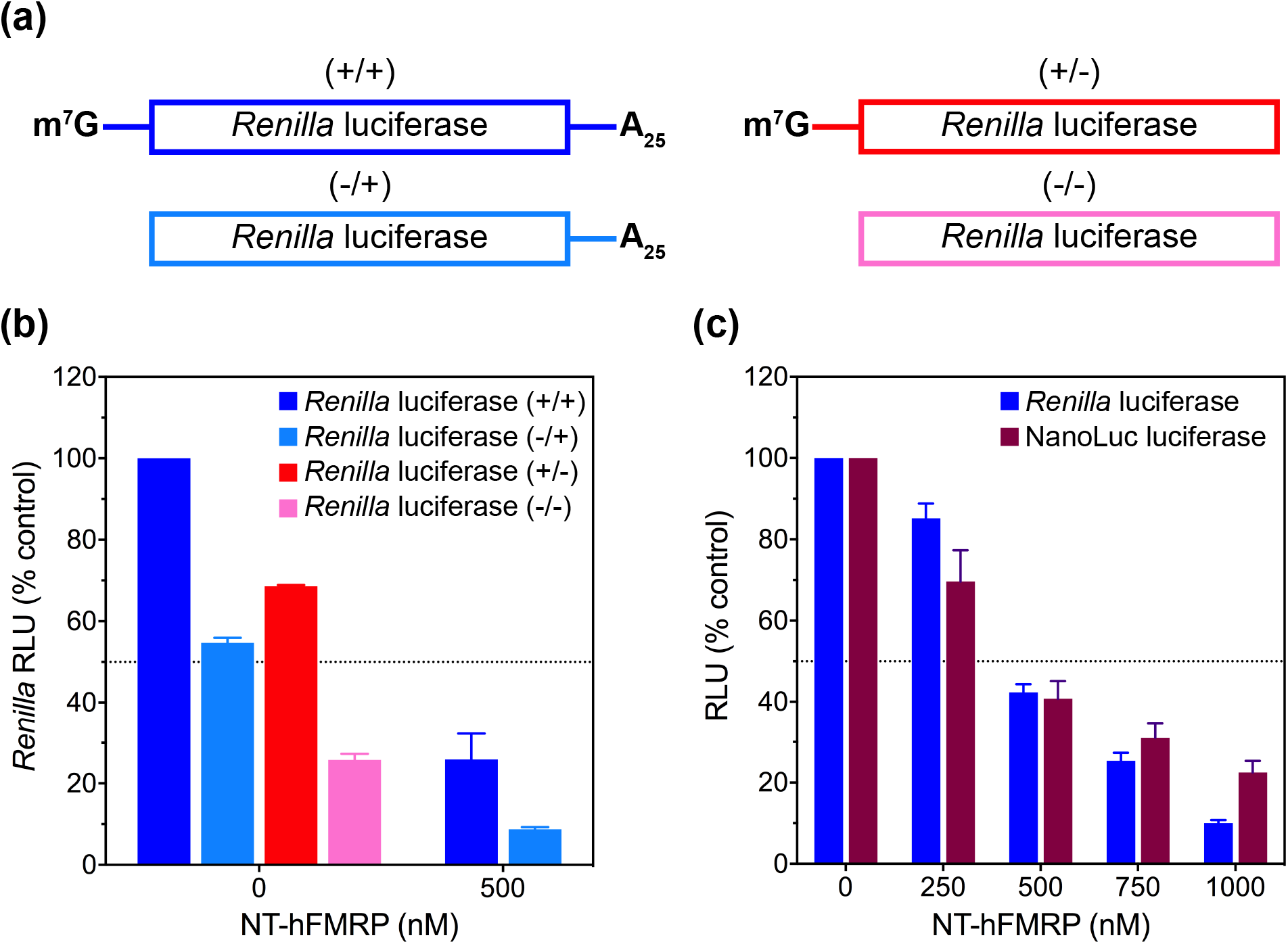
Human FMRP inhibits translation of different mRNAs in RRL. (a) Diagram of *Renilla* luciferase (RLuc) mRNAs containing: both 5’-m^7^G cap and 3’-polyA25 tail (+/+, blue), only 3’-polyA25 tail (-/+, cyan), only 5’-m^7^G cap (+/-, red), or neither 5’-m^7^G cap nor 3’-polyA25 tail (-/-, pink). (b) Relative luminescence units (RLU) from 10 nM of the four aforementioned *Renilla* luciferase mRNAs, and upon the addition of 500 nM NT-hFMRP, all as a percentage of the (+/+) RLuc mRNA. (c) RLU from 10 nM *Renilla* luciferase (blue) and 10 nM NanoLuc luciferase (maroon) upon the addition of 0-1000 nM NT-hFMRP as a percentage of the control containing no NT-hFMRP. The standard deviations from three experiments are shown.

However, it was unclear whether FMRP specifically inhibits translation of RLuc mRNA because it contains a feature that makes it a target mRNA, or if FMRP would have a similar effect on the translation of different mRNAs. Therefore, we compared the effect of NT-hFMRP on the translation of *Renilla* luciferase (RLuc) and NanoLuc luciferase (NLuc) reporter mRNAs. Surprisingly, we observed nearly identical effects on both mRNAs, suggesting FMRP indiscriminately inhibits translation through a common mechanism **(Figure 2c)**.

### FMRP inhibits the elongation step of translation

We observed that hFMRP equally inhibited the translation of 5’-capped and uncapped RLuc mRNAs **(Figure 2b)**. Therefore, we sought to test hFMRP inhibition of canonical capped-and-tailed RLuc mRNA against internal ribosome entry site (IRES)-driven mRNAs. We generated RLuc mRNAs driven by a representative IRES from the four different classes of IRESes: poliovirus (PV-RLuc), encephalomyocarditis virus (EMCV-RLuc), hepatitis C virus (HCV-RLuc), and cricket paralysis virus (CrPV-RLuc) [49]. Unfortunately, the PV-RLuc and HCV-RLuc mRNAs did not translate efficiently enough for IVT studies, so we proceeded with the EMCV-RLuc and CrPV-RLuc mRNAs at a fixed 10 nanomolar concentration for comparison against 10 nanomolar canonical RLuc **(Figure 3a)**. While one may expect more protein synthesis from having more mRNAs in the IVT reaction, too much of the EMCV IRES-driven RLuc mRNA actually decreases translation efficiency [50]. Consistent with our previous observation, we saw FMRP inhibits the translation of CrPV-RLuc as efficiently as the canonical RLuc, suggesting that FMRP indeed inhibits translation of RLuc exclusively at the elongation step because CrPV IRES mRNA does not require any of the initiation factors to bind to the ribosome and initiate protein synthesis [51–54]. Inhibition of EMCV-RLuc was slightly milder compared to canonical RLuc and CrPV-RLuc **(Figure 3b)**.

**Figure 3.**
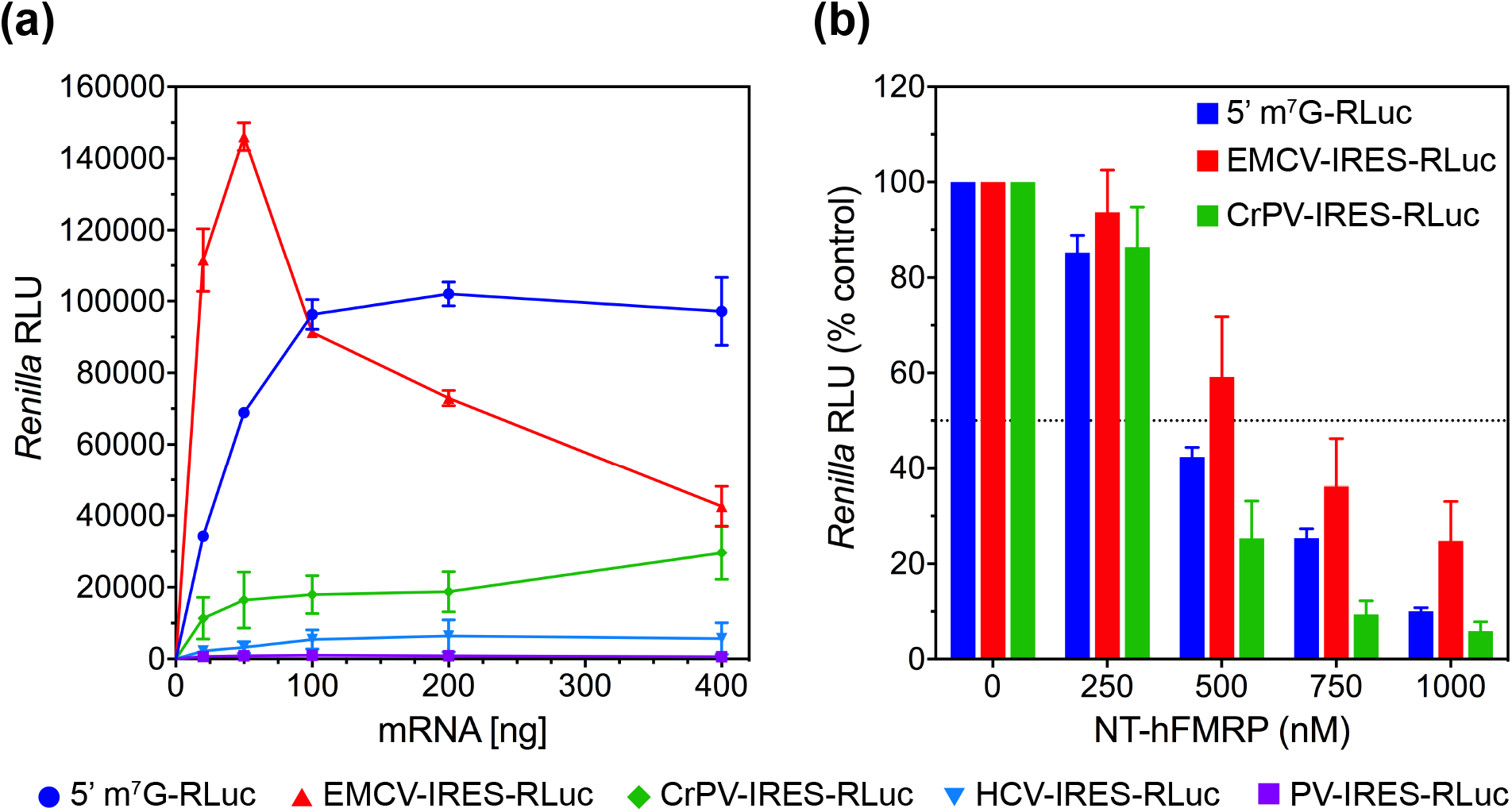
FMRP inhibits translation of canonical and IRES-driven *Renilla* luciferase mRNAs. (a) Relative luminescence units (RLU) from *Renilla* luciferase upon the addition of 0-400 ng of the indicated mRNAs in RRL. (b) RLU from canonical *Renilla* luciferase (blue), EMCV IRES-driven *Renilla* luciferase (red), and CrPV IRES-driven *Renilla* luciferase (green) upon the addition of 0-1000 nM NT-hFMRP as a percentage of the control containing no NT-hFMRP. The standard deviations from three experiments are shown.

### The RGG domain and CTD are sufficient to inhibit translation

To identify which regions and RNA-binding domains of hFMRP contribute to translation inhibition, we measured the effects of the GST-tagged KH0-KH1-KH2 region (GST-KH0-KH1-KH2), GST-tagged RGG-CTD region (GST-hFMRP RGG), and a GST-tagged mutant SGG-CTD region (GST-hFMRP SGG) on RLuc mRNA translation. We found that GST-hFMRP RGG was sufficient to inhibit the translation of RLuc **(Figure 4a)**. In contrast, GST-KH0-KH1-KH2 did not inhibit translation and had a similar effect as the mutant GST-hFMRP SGG in which the RNA G-quadruplex (GQ)-binding arginines spanning the RGG domain and CTD of FMRP were all mutated to serines **(Figure 4a)**. Furthermore, we tested whether the RNA GQ-binding RGG domain was sufficient to inhibit translation. We used the 18 amino acid RGG peptide previously found to bind the sc1 GQ-forming RNA and observed it was insufficient to inhibit translation, suggesting the CTD of hFMRP plays an essential role in translation inhibition **(Figure 4a)** [23,55]. Finally, the GST-hFMRP RGG is not as efficient at inhibiting RLuc translation compared to NT-hFMRP. This suggests there may indeed be a cooperative effect between the C-terminus of hFMRP, containing the RGG and CTD, the KH1 and KH2 domains, and the unstructured region between the KH2 and RGG domains. Altogether, it is possible the RGG and CTD are critical for the initial binding interaction as an anchor while the other regions of FMRP further stabilize that interaction.

**Figure 4.**
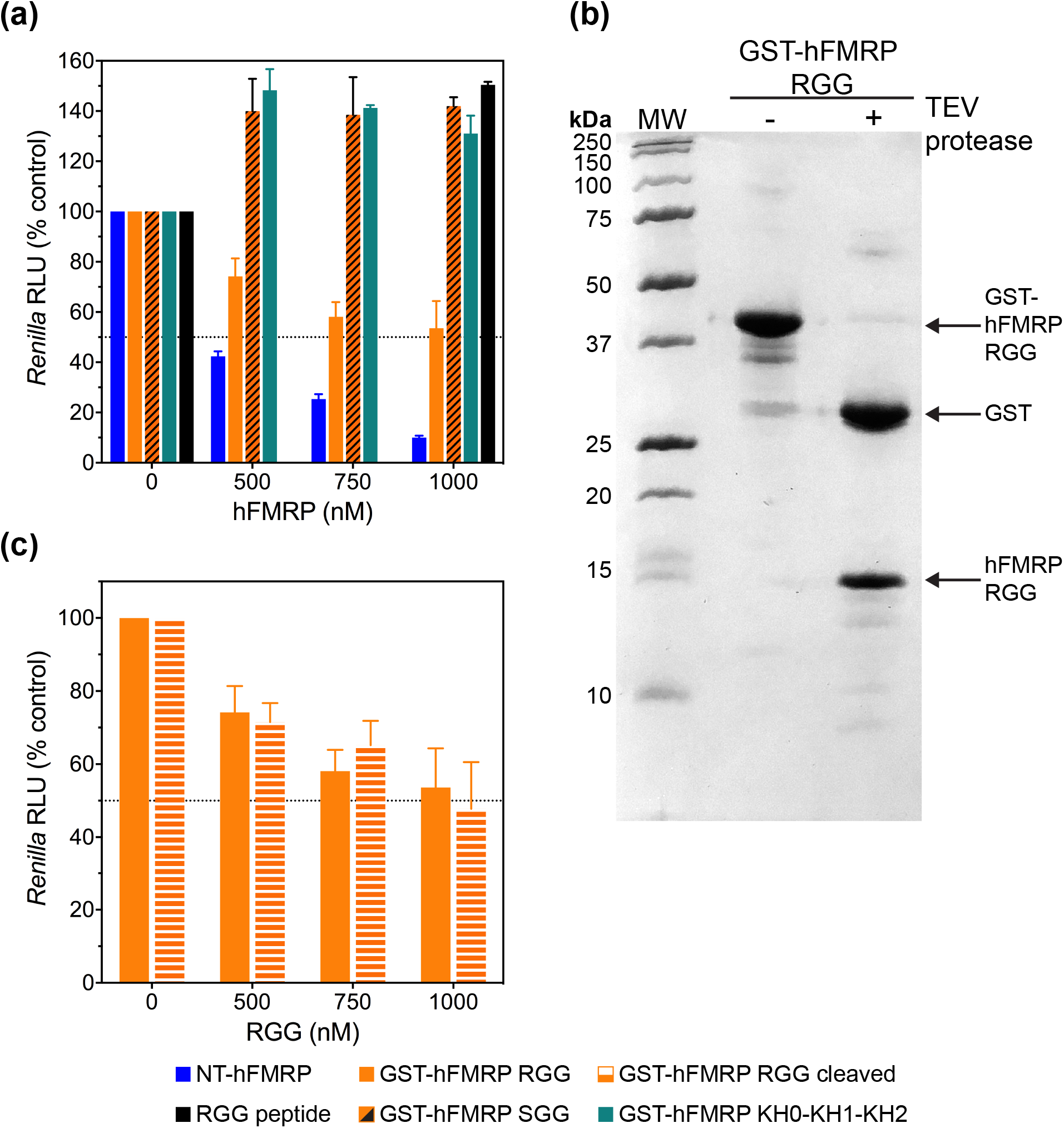
The RGG domain and CTD of FMRP are required for inhibition of translation. (a) Relative luminescence units (RLU) from *Renilla* luciferase upon the addition of 0-1000 nM NT-hFMRP (blue), GST-hFMRP RGG (orange), GST-hFMRP SGG (orange with black diagonal stripes), GST-hFMRP KH0-KH1-KH2 domains (teal), and RGG peptide (black) in RRL. (b) 12% SDS-PAGE of GST-hFMRP RGG before (-TEV protease) and after (+ TEV protease) cleaving with TEV protease. (c) RLU from *Renilla* luciferase upon the addition of 0-1000 nM GST-hFMRP RGG (orange) and GST-hFMRP RGG precleaved using TEV protease (orange with white horizontal stripes) in RRL. The standard deviations from three experiments are shown.

To determine if the GST tag is involved in translation inhibition by the RGG-CTD region, we cleaved the GST-hFMRP RGG with TEV protease to separate the GST tag and RGG-CTD fragments before IVT **(Figure 4b)**. We observed no significant difference in translation inhibition between the intact GST-hFMRP RGG fusion protein and the cleaved protein, showing that the GST protein is not involved in translation inhibition by the RGG-CTD fragment **(Figure 4c)**.

### FMRP binding to the ribosome correlates with inhibition of translation

Based on past findings and our observations that FMRP inhibits translation at the elongation step and the RGG-CTD region is sufficient for inhibition, we hypothesized the RGG-CTD might be serving as the anchor during direct binding of FMRP to the ribosome. Two recent studies found specific rRNA expansion segments in the ribosomes of higher eukaryotes contain G-quadruplex structures [37,38]. Together with our recent study showing the RGG domain of hFMRP can bind even a minimal G-quadruplex structure, we investigated whether the RGG-CTD region of hFMRP could be binding directly to a G-quadruplex on the ribosome to inhibit translation [24]. To determine whether binding to the ribosome is correlated with translation inhibition, we tested the binding of the human 80S ribosome to NT-hFMRP, GST-hFMRP RGG, and GST-hFMRP KH0-KH1-KH2 by sucrose cushion co-sedimentation. We observed both NT-hFMRP and GST-hFMRP RGG bind directly to the ribosome, while GST-hFMRP KH0-KH1-KH2 does not bind **(Figure 5)**. Our results suggest that the direct binding of FMRP to the ribosome is crucial for inhibition of translation.

**Figure 5.**
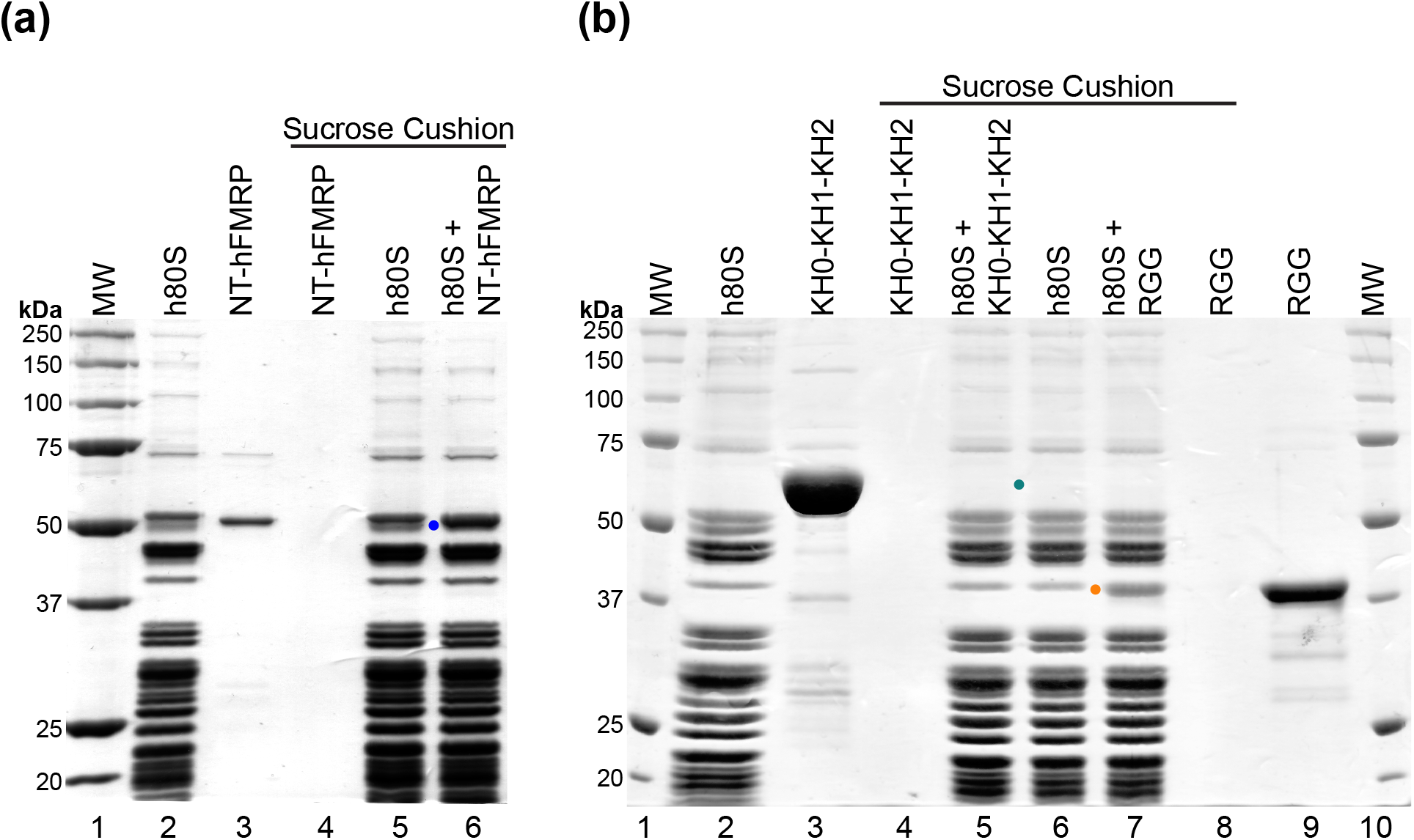
NT-hFMRP and GST-hFMRP RGG bind directly to the human 80S ribosome. (a) A representative 10% SDS-PAGE of 40% sucrose co-sedimentation assay showing NT-hFMRP (blue dot) co-sediments with human 80S ribosomes (h80S). Samples in lanes 4 to 6 were sedimented through the sucrose cushion. Lanes: 1, molecular weight ladder; 2, input h80S; 3, input NT-hFMRP; 4, NT-hFMRP in pellet; 5, h80S in pellet; 6, h80S+NT-hFMRP complex in pellet. NT-hFMRP migrated close to a ribosomal protein; however, quantification showed that the band intensity of the NT-hFMRP and the ribosomal protein in lane 6 increased by 25% compared to the same ribosomal protein band in lane 5. (b) A representative 10% SDS-PAGE of 40% sucrose co-sedimentation assay showing GST-hFMRP RGG (orange dot) co-sediments with human 80S ribosomes, but GST-hFMRP KH0-KH1-KH2 (teal dot) does not. Samples in lanes 4 to 8 were sedimented through the sucrose cushion. Lanes: 1, molecular weight ladder; 2, input h80S; 3, input KH0-KH1-KH2; 4, KH0-KH1-KH2 in pellet; 5, h80S+KH0-KH1-KH2 in pellet; 6, h80S in pellet; 7, h80S+GST-RGG in pellet; 8, GST-RGG in pellet; 9, input GST-RGG; 10, molecular weight ladder. GST-RGG migrated close to a ribosomal protein; however, quantification showed that the band intensity of the GST-RGG and the ribosomal protein in lane 7 increased by 30% compared to the same ribosomal protein band in lane 6.

## Discussion

Previous studies have shown that FMRP inhibits a variety of natural and artificial reporter mRNAs suggesting that these mRNAs may have a common RNA motif that is recognized by FMRP [56–61]. Alternatively, FMRP may globally inhibit translation by repressing the function of a specific translation factor or the ribosome. Our IVT studies in RRL lead us to conclude that human FMRP inhibits mRNAs indiscriminately and to similar degrees when concentrations of two different mRNAs of different lengths are standardized. This suggests FMRP does not necessarily regulate the translation of only a subset of target mRNAs, but can also regulate mRNAs globally through a common upstream translation mechanism independent of sequence or structure features in the mRNA.

FMRP also potently inhibits the translation of RLuc mRNA independent of the 5’ cap, showing that potent inhibition can occur without FMRP interacting with the initiation machinery. To confirm our findings, we used CrPV IRES, which binds directly to the ribosomal P site to initiate translation without the need for any initiation factors [51–54]. FMRP inhibits CrPV-RLuc just as potently as the canonical RLuc, supporting the hypothesis that FMRP inhibits at the elongation step regardless of the initiation rate and initiation machinery. We observe milder inhibition of EMCV-RLuc by FMRP, which may suggest FMRP’s ability to inhibit translation is linked to factors beyond translation elongation. A recent study looked into the differences between canonical and EMCV-driven translation and found EMCV IRESes are indistinguishable from canonical 5’ UTRs in terms of initiation and elongation rates. However, the EMCV IRES is less efficient at recruiting ribosomes than the 5’ cap machinery [62]. Ribosome recruitment and availability, together with elongation, seem to be major factors for FMRP inhibition of translation.

Upon dissecting FMRP, we determined the arginine-rich RGG domain and CTD are the primary drivers of translation inhibition, while the region spanning the putative RNA-binding KH0 to KH2 domains alone cannot inhibit translation. Although the RGG-CTD region may be sufficient for mild translation inhibition, potent inhibition requires additional features found in NT-hFMRP, such as the unstructured region between the KH2 and RGG domains. Based on our ribosome-binding study, it is possible that the KH domains cannot inhibit translation alone because they cannot form a stable interaction with the ribosome. Perhaps the KH domains cooperate with the RGG-CTD to stabilize the initial interaction between the RGG-CTD and the ribosome to enhance translation inhibition.

Overall our studies suggest hFMRP can globally inhibit mRNA translation by binding directly to ribosomes through its RGG domain and CTD tail to (1) effectively decrease the population of active ribosomes available to translate mRNAs, and (2) stall actively translating ribosomes at the elongation step. It remains to be seen whether RREs, such as G-quadruplexes, within the mRNAs can alter translation regulation by FMRP.

## Materials and Methods

### Protein expression and purification

The human FMR1 isoform 1 gene was assembled from *E. coli* codon-optimized gene blocks (IDT). The gene encoding the full-length isoform 1 human FMRP (hFMRP) spanning residues E2-P632 was subcloned into the LIC expression vector pMCSG7 (DNASU plasmid repository) with the 5’ TEV cleavage site deleted, conferring a N-terminal hexahistidine tag. The N-terminus truncated human FMRP (NT-hFMRP) spanning residues R218-P632 was subcloned into the LIC expression vector pMCSG7, conferring a N-terminal hexahistidine tag with the 5’ TEV cleavage site deleted. The human FMRP RGG domain (GST-hFMRP RGG) spanning residues G531-P632 was subcloned into the LIC expression vector pMCSG10 (DNASU) which confers a N-terminal hexahistidine-GST fusion tag that is cleavable using TEV protease. The mutant SGG domain (GST-hFMRP SGG) was assembled from an *E. coli* codon-optimized gene block where all 16 arginine residues of the RGG region were mutated to serines. The resulting SGG fragment was subcloned into the LIC expression vector pMCSG10. The human FMRP tandem KH0-KH1-KH2 domains (GST-hFMRP KH0-KH1-KH2) spanning residues F126-K425 was also subcloned into the LIC expression vector pMCSG10.

The NT-hFMRP, GST-hFMRP RGG, GST-hFMRP SGG, and GST-hFMRP KH0-KH1-KH2 expression plasmids were all transformed in *E. coli* BL21(DE3) cells (Novagen). NT-hFMRP was purified using Ni-NTA affinity chromatography (Qiagen). GST-hFMRP RGG, GST-hFMRP SGG, and GST-hFMRP KH0-KH1-KH2 were purified using glutathione affinity chromatography (GE Healthcare). All proteins were further purified using Superdex 75 16/60 or Superdex 200 16/60 (GE Healthcare) gel filtration chromatography and stored in their respective running buffers. Gel filtration buffer composition varied between FMRP constructs. NT-hFMRP gel filtration running buffer contained 25 mM Tris pH 7.4, 150 mM KCl and 1 mM DTT. GST-hFMRP RGG and GST-hFMRP SGG were purified in gel filtration running buffer containing 50 mM Tris pH 7.5 and 1 mM DTT. GST-hFMRP KH0-KH1-KH2 was purified in gel filtration running buffer containing 50 mM Tris pH 7.4, 150 mM NaCl and 1 mM DTT.

### Preparation of reporter mRNAs

The gene encoding the *Renilla* luciferase (RLuc) protein was modified with an poly(A)25 insertion at the 3’ end of the gene within the pRL-null vector (Promega). The gene encoding the NanoLuc luciferase was transferred from the pNL1.1 vector (Promega) into the pRL-null vector using 5’ NheI and 3’ XbaI restriction enzymes, then a poly(A)25 was inserted at the 3’ end of the NanoLuc gene. This generated RLuc and NanoLuc mRNAs with uniform 3’ poly(A)25 tails to bypass the need for post-transcriptional 3’ polyadenylation. The PV IRES-, EMCV IRES-, HCV IRES-, and CrPV IRES-driven RLuc genes were subcloned into the EMCV-TurboGFP parental vector pT7CFE1 (ThermoFisher Scientific) in two steps. In the first step, the TurboGFP gene was replaced with the RLuc gene using 5’ NdeI and 3’ XhoI restriction enzymes to generate the EMCV-RLuc plasmid. In the second step, the EMCV IRES was replaced with either the PV, HCV, or CrPV IRES using 5’ ApaI and 3’ NdeI restriction enzymes to generate the PV-RLuc, HCV-RLuc, and CrPV-RLuc plasmids, respectively. The four IRES-driven RLuc genes contain a 3’ poly(A)_30_ sequence from the pT7CFE1 vector, which resulted in the synthesis of mRNAs with uniform 3’ poly(A)_30_ tails.

All 6 mRNAs were synthesized by run-off in vitro transcription using T7 RNA polymerase at 37°C for 4-6 hours. The RLuc template DNA was prepared by linearizing the vector using XbaI, while the NanoLuc and IRES-driven template DNAs were prepared by PCR amplification. The transcription reactions were treated with RQ1 DNase (Promega) at 37°C for 30 minutes to remove the DNA templates. Each transcription reaction was cleaned up in two steps, first by chloroform extraction and ethanol precipitation followed by Monarch RNA cleanup kit (NEB).

The RLuc and NanoLuc mRNAs were post-transcriptionally 5’ capped with 7-methylguanosine (m^7^G) using a homemade Vaccinia capping enzyme (REF). The mRNAs were first heated for 5 minutes at 65°C and then put on ice for 5 minutes. Capping was carried out for 2 hours at 37°C in 50 μL reactions with 1X capping buffer (50 mM Tris pH 8, 5 mM KCl, 1 mM MgCl2, and 1 mM DTT), 0.5 mM GTP, 0.1 mM SAM, and 4 μL homemade Vaccinia capping enzyme; the amount of mRNA substrate to be modified was limited to maintain a large stoichiometric excess of both GTP and SAM to ensure all mRNA molecules were capped. The 5’ m^7^G-capped mRNAs were then cleaned up using the Monarch RNA cleanup kit.

### In vitro translation assays

The rabbit reticulocyte lysate was prepared following the method described by (1) Feng and Shao, and (2) Sharma *et al* [63,64]. In vitro translation (IVT) reactions in rabbit reticulocyte lysate (RRL) were carried out in a final 20 mL volume at 30°C for 90 minutes. First, in a 10 mL binding reaction, 24 mM HEPES pH 7.5, 100 mM potassium acetate, 20 nM mRNA, and 500-2000 nM protein were combined and incubated for 15 minutes at room temperature; the control sample contained the protein storage buffer equivalent to the highest volume of protein used in the titration (maximum 2 mL). Then 10 mL RRL was added to make the final 20 mL IVT reaction. After 90 minutes at 30°C, the samples were brought to room temperature for 5-10 minutes while 2 mL of 30 mM coelenterazine (Promega), or 15 mL of 1-to-50 diluted NanoLuc substrate (Promega), was pipetted into a 384-well white flat bottom plate (Greiner). In the case of IVT reactions containing RLuc mRNAs, 18 mL of the 20 mL sample was mixed with 2 mL coelenterazine, and luminescence readings were taken immediately on a Tecan Spark plate reader. In the case of IVT reactions containing NanoLuc mRNAs, 15 mL of the 20 mL sample was mixed with 15 mL of 1-to-50 diluted NanoLuc substrate, and luminescence readings were taken immediately on a Tecan Spark plate reader. Luminescence readings of samples in each titration were normalized to the signal of their respective control sample.

For the GST-hFMRP RGG cleavage reactions with TEV protease, GST-hFMRP RGG fusion protein was combined with either TEV protease or storage buffer (50 mM Tris pH 7.4 and 1 mM DTT) at a 4:1 ratio for a final 20 mM GST-hFMRP RGG. The cleavage reaction was carried out for 2 hours and 30 minutes at room temperature with gentle agitation, after which the protein samples were serially diluted in storage buffer for the IVT assay.

### Human 80S ribosome binding assay

In a 20 mL binding reaction, 0.8 mM human 80S ribosomes and 7.2 mM protein were incubated with 50 mM Tris acetate pH 7.7, additional potassium acetate (final 50 mM K^+^), additional magnesium acetate (final 3.8 mM Mg^2+^), 10 mM DTT, and 30 mg/mL total tRNA from *E. coli*. The 20 mL reactions were incubated for 15 minutes at room temperature, placed on ice for 2 minutes, and then carefully pipetted onto a pre-cooled 200 mL 40% (w/w) sucrose cushions reconstituted in RBB (50 mM Tris acetate pH 7.7, 50 mM potassium acetate, 5 mM magnesium acetate, and 10 mM DTT). The sample was centrifuged at 4°C for 1 hour and 40 minutes at 182,753 x g (38,000 rpm in a Beckman Coulter Type 42.2 Ti rotor).

The supernatant was carefully removed then 16 mL RBB was gently pipetted onto the ribosome pellet and incubated covered at 4°C for 30 minutes. The 16 mL sample was carefully mixed by gentle pipetting in place and transferred to a clean tube containing 4 mL 5X SDS sample loading solution. 10 mL of the 20 mL gel sample was analyzed by 10% SDS-PAGE and stained with Coomassie brilliant blue.

